# Predicting Decision-Making Time for Diagnosis over NGS Cycles: An Interpretable Machine Learning Approach

**DOI:** 10.1101/2023.03.07.530760

**Authors:** Athar Khodabakhsh, Tobias P. Loka, Sébastien Boutin, Dennis Nurjadi, Bernhard Y. Renard

## Abstract

**Motivation:** Genome sequencing processes are commonly followed by computational analysis in medical diagnosis. The analyses are generally performed once the sequencing process has finished. However, in time-critical applications, it is crucial to start diagnosis once sufficient evidence has been accumulated. This research aims to define a proof-of-principle for predicting earlier time for decision-making using a machine learning approach. The method is evaluated on Illumina sequencing cycles for pathogen diagnosis.

**Results:** We utilized a Long-Short Term Memory (LSTM) approach to make predictions for the early decision-making time in time-critical clinical applications. We modeled the (meta-)information obtained from NGS intermediate cycles to investigate whether there are any changes to expect in the remaining sequencing cycles. We tested our model on different patient datasets, resulting in high accuracy of over 98%, indicating the model is independent of a dataset. Furthermore, we can save several hours of turnaround time by using the early prediction results. We used the SHapley Additive exPlanations (SHAP) framework for the interpretation and assessment of the LSTM classifier.

**Availability:** The source code is available at https://gitlab.com/dacs-hpi/ngs-biclass.

## 1 Introduction

Highly accurate genomic information extracted by next-generation sequencing (NGS) provides immense insights about biological systems. The development of second and third generation approaches has revolutionized DNA sequencing. Furthermore, modern DNA sequencing techniques have led to improvements in disease-related diagnosis [Hong *et al*., 2014]. Consequently, genome sequencing has matured and is arriving in clinical applications [Andrusch *et al*., 2018]. Several areas of medical diagnosis have already benefited from genome sequencing, such as the diagnosis of genetic diseases in very ill infants [Willig *et al*., 2015], the genotype-guided choice of chemotherapy [Tran *et al*., 2013], and pathogen diagnostic [Naccache *et al*., 2014]. In such situations, delayed decisions may lead to severe medical conditions [Miller *et al*., 2015]. The wealth of information obtained by sequencing is unmatched, but for some applications (e.g., pathogen diagnosis [Mongan *et al*., 2020] and in particular, the detection of sepsis [Husabø *et al*., 2020]), even hours in overall turnaround times can have an impact. In conventional NGS-based approaches, analysis procedures start once sequencing has finished, which results in a high turnaround time from sample arrival to identification and diagnosis. Increasingly, rapid procedures are available for sequencing. For second-generation platforms (such as Illumina), tools are available to analyze data during runtime (e.g., LiveKraken [Tausch *et al*., 2018] and Hilive2 [Loka *et al*., 2019]), or rapidly after the end of the run (e.g., [Miller *et al*., 2015] and [Linder *et al*., 2017]). For third-generation sequencing, particularly Nanopore, reads are sequentially acquired and analyzed [Mikheyev *et al*., 2014]. Adaptive sequencing methods such as MinKNOW [Sim *et al*., 2019] [Runtuwene *et al*., 2019], Readfish [Payne *et al*., 2021], and ReadBouncer [Ulrich *et al*., 2022] can help to enrich for targeted regions and thereby allow faster diagnosis.

The challenge of efficient clinical decision-making is explored in many diagnostic procedures; however, reducing the medical decisionmaking time is less studied in genome diagnosis. For instance, in the treatment of advanced tumors by radiation therapy (RT), [Brahme, 2000] developed an optimization approach to improve treatment time, treatment fractions, and accuracy of dose response. [Eikelder *et al*., 2019] proposed an adaptive strategy based on mid-treatment biomarker information to adjust the treatment length of an RT. While accurate results have the highest priority in clinical applications, time is the second crucial factor in allowing optimal treatment decisions as early as possible. This is reflected in a speed-accuracy trade-off in decision-making [Forstmann *et al*., 2016]. The early information can lead to fast decisions, but it can sacrifice accuracy. In contrast, postponing the decision until all evidence is gathered can be very late, i.e., diagnosis under time pressure for a patient in a clinical setting. Therefore, mechanisms are required for efficient decision-making once enough evidence has been accumulated [Drugowitsch *et al*., 2019].

The motivation of this work is a data-driven analysis of sequencing (meta-)information to predict the earliest time for diagnosis, specifically when more sequencing is not leading to more relevant information and, thereby, to other decision options. The *proof-of-principle* that we proposed is based on Illumina sequencing for pathogen diagnosis, but the ideas are more generally applicable in clinical diagnosis. Our approach is to trigger an early decision by learning and making predictions with respect to the accumulated information at earlier time points. We propose to utilize a supervised machine learning approach using a long-short term memory (LSTM) [Hochreiter *et al*., 1997] neural network for the prediction and decision-making challenge observed in NGS-based diagnosis. Recurrent neural networks (RNN) are robust neural networks for modeling sequence data such as natural language processing or time-series [Lipton *et al*., 2015]. The functionality of RNN is iterations over time steps of a sequence while internal states are maintained for observed sequences. LSTMs are RNNs that can learn order dependence in sequence prediction problems. The advantage of the LSTM is that it overcomes the vanishing gradient problem [Hochreiter, 1998] of RNNs and is effective in capturing long-term temporal dependencies [Bengio *et al*., 1994]. To address the speed-accuracy trade-off, we conceptualize the accumulated information with time-series analysis methods by defining a sliding window model [Braverman *et al*., 2009] for making a prediction based on the available information from the intermediate stages. Furthermore, understanding the reasons behind predictions is crucial for assessing trust, which is fundamental in taking action based on a prediction, or in predictive model deployment [Ribeiro *et al*., 2016]. There are several approaches to interpreting machine learning models, such as local interpretable model-agnostic explanations (LIME) [Ribeiro *et al*., 2016] and SHapley Additive exPlanations (SHAP). SHAP is a game-theoretic approach to explain the output of any machine learning model [Lundberg *et al*., 2017]. We relied on the SHAP framework to assess the model outputs.

## 2 Method

Our approach for predicting earlier time for diagnosis is to use the output of real-time NGS sequence analysis to learn properties from incomplete fragments of genome reads. Figure 1 illustrates this conceptual idea. The high turnaround time challenge is first addressed by real-time analysis as shown in Figure 1, where post-hoc processing is not required after the sequencing run has finished. Secondly, for the time-critical aspect of diagnosis, we aim to push the analysis further to an earlier time point by using prediction results to have additional time saving in parallel to real-time analysis. Ideally, an earlier time point for diagnosis is possible by using predictive methods. Thus, the information extracted from the intermediate analysis is used for building a predictive model. The underlying idea is to use quality data of early genomic read alignments to predict whether changes in the alignments are likely to occur. Readmapping quality may change over time while the length of genome reads increases in each cycle.

**Fig. 1.**
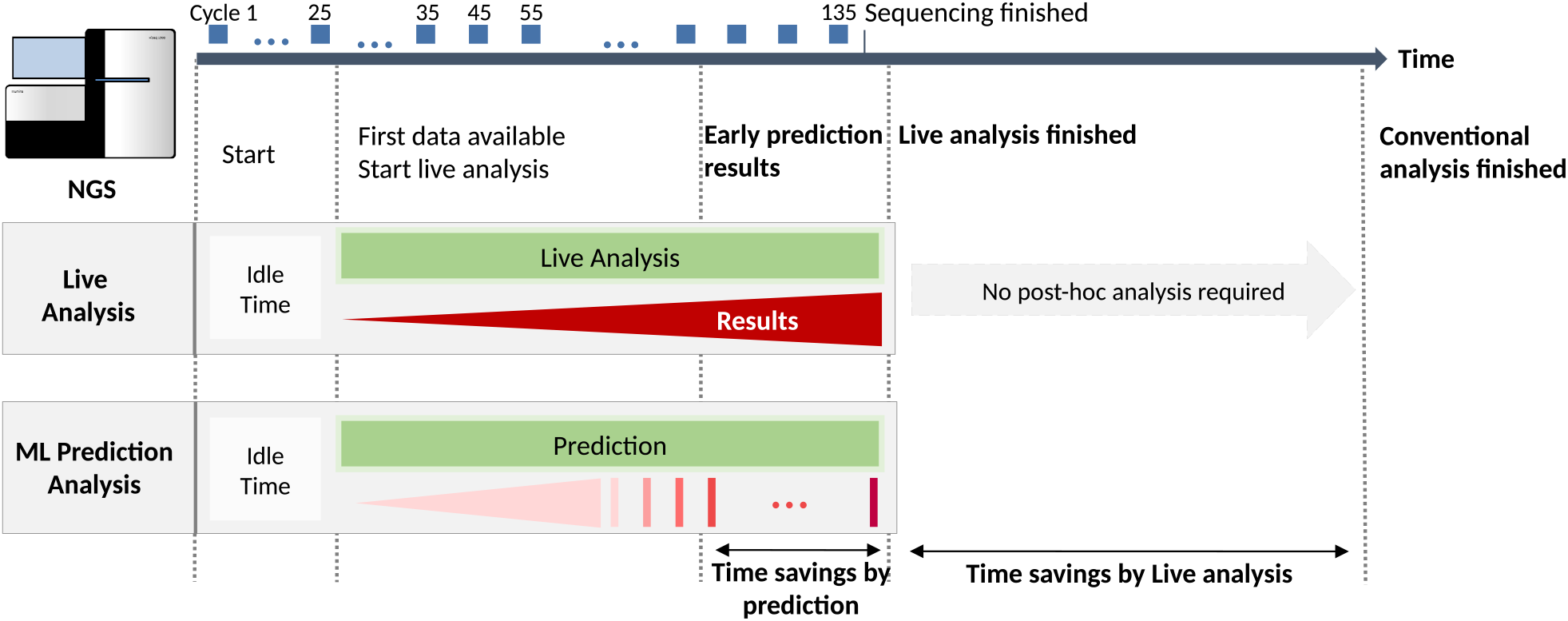
The conceptual idea is to reduce time-to-decision in time-critical applications. The intermediate results obtained from the NGS cycles are analyzed by a real-time read-mapping algorithm such as HiLive2 [Loka *et al*., 2019]. The alignment quality data are used to train an LSTM binary classifier and predict changes in future cycles. The prediction performances are monitored to recognize the earliest time for early diagnosis. Earlier decisions can be vital in diagnosis under time pressure for patients in a clinical setting.

In our experiments, we used the read quality (meta-)information to predict if a read identified and matched to a pathogen will continue with the same database hit, until the sequencing will have ended. In some cases, initial matches on very short sequence stretches may be discarded later as the number of mismatches increases. In other cases, short sequence stretches from early cycles may not be specific enough to match a particular sequence, and only as the read extends in length the read alignment becomes feasible. We propose to use a data-driven approach by training a binary classifier using an LSTM model to predict alignment changes. A true result means that a change in an alignment is less likely to happen. In contrast, a false output indicates that the alignment has an alteration possibility in future cycles. The benefit of binary classification is an explicit indication about a genome read to expect in the remaining sequencing cycles. We modeled the data by sliding window model using LSTM iteratively over intermediate cycles. The performances of the LSTM models are monitored over time to recognize the best-performing window in any of the intermediate stages. Furthermore, to interpret the contributions of genomic read alignment (meta-)information on model output, we assessed the LSTM classifier with the SHAP framework, which explains the model’s output with local explanations [Lundberg *et al*., 2020] using classic Shapley values [Shapley, 1953]. We used the SHAP framework because it has the advantage of decomposing final predictions into the additive contribution of each attribute, which is not provided in LIME. The interpretable machine learning approach in this paper consists of the following steps:

- Data preprocessing: read alignment (meta-)information obtained from real-time NGS-based analysis cycles is cleaned, normalized, and labeled to prepare a sequential data format.
- Binary classifier: an LSTM neural network is utilized as a binary classifier of reads to predict changes likely to occur during intermediate cycles.
- Evaluation: classification performances of the models are evaluated on test data; from the same sample and four different patient samples.
- Decision-making time: the classification performances are monitored over eight sliding windows for estimation of earlier time point for decision-making.
- Interpretable machine learning: Shapley values are extracted to assess the black-box neural network and explain the LSTM binary classifier’s outputs.

### 2.1 Preprocessing of Sequential Data

#### 2.1.1 Intermediate Cycles

NGS sequencer generates intermediate batches of genome reads at evenly spaced time points known as sequencing “*cycles*.” Each cycle consists of (meta-)information of sequenced nucleotides for fragments of DNA known as “*reads*.” The length of a genome sequence in a cycle is equal to the number of respective cycle. For instance, in cycle 45, the sequences contain 45 base pairs. During analysis, the genome reads from each cycle are aligned to genome references resulting in an assignment. Therefore, the new sequence (meta-)information obtained in each sequencing cycle can be used in real-time to extend the previous state of the genome analysis. Each read has several features that represent the alignment quality. Differences in the mapping between the sequence in a read and the reference genome are scored with penalties; thus, we get lower scores for higher discrepancies and errors. The (meta-)information used in preprocessing in this study is listed below. However, other information, such as the genome sequence itself, CIGAR string, etc., could be added to the analysis, which is not the focus of this study.

Read-ID: is an identifier that tracks genomic reads in cycles. A read assigned to a reference genome during sequencing will appear with the same ID at all the cycles. It is worth noting that the reads keep their ID whether they appeared or were discarded in intermediate cycles.
MAPQ: represents the mapping quality. Higher MAPQ values imply the read’s alignment position to the reference genome is accurate [SAM Tools, 2021].
AS: represents another mapping score known as alignment score. If the sequence is identical to the sequence in the reference database, the AS value will be 0. A penalty is applied for each error/difference; thus, the lower the score, the higher the number of errors in the read alignment.
Pathogen-ID: when a read has been assigned to a reference genome, the identified “*genus*” will be stored as a feature in each cycle. In this study, change of reference assignment is only studied at the “*genus*” level. While the alignment itself can be more specific, we do not aim to predict changes on the sub-genus level since only in very rare cases this can relate to changes in pathogenicity [Deneke *et al*., 2017] in gram-negative bacteria.

The alignment quality data is collected from consecutive cycles in time order. We used this (meta-)information as features for preparing a sequential data structure. We used MAPQ, AS, and Pathogen-ID features in this study. Read-ID is a (meta-)information used to track sequence fragments and interconnecting information, which is not considered a feature for modeling since it is merely an identifier. After the dataset has been prepared, the Read-IDs are eliminated.

#### 2.1.2 Sliding Window Model

In principle, a sliding window model is applied for streams of data where only the most “recent” elements stay active, and the rest are discarded [Braverman *et al*., 2009]. We used a fixed-time window model where data arrive in time order, and just the recent *n* elements are captured. In this study, as a batch of new alignment quality data arrives from each intermediate cycle, they are added consecutively in time order *t, t* + *i,t* + 2*i*,.., where *t* is stating cycle time, and *i* is the time-step. We process three consecutive cycles in sequential order. This structure is depicted in Figure 2, where information from the most recent three cycles is used. According to our experiments, we could perform an efficient sequential data analysis by building an LSTM classifier with *n* = 3 and *i* = 10. A larger *n* leads to a higher time delay which is not appropriate in time-critical analysis, and lower window sizes showed insufficient accuracy of results.

**Fig. 2.**
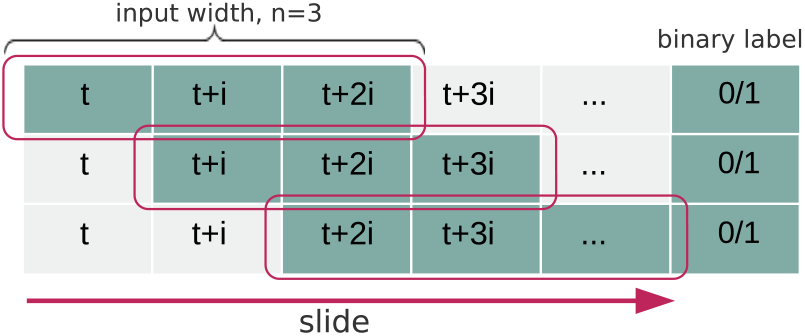
Sliding window model applied in preprocessing steps for preparing the sequential data. We set the window size to *n* = 3 and the time-step to *i* = 10.

### 2.2 Binary Classification

The early genomic read alignment quality information is used to predict whether changes in the alignments are likely to occur by binary classification. The reads classified as true have a higher probability of occurring in future cycles; in contrast, false reads are less likely to appear. To accomplish classification on sequential data, the sliding window model shown in Figure 2 is used as input for the LSTM classifier. Each window consists of quality data of genomic read alignment of respective cycles. For instance, a window at 45-55-65 contains MAPQ, Pathogen-ID, and AS of cycles 45, 55, and 65 consecutively. In each window, reads are labeled with true or false, where the binary classes are conceptually defined as follows:

**true** class: if a read has been assigned to the same Pathogen-ID in all three cycles, we consider it as a true read.
**false** class: if a read’s Pathogen-ID changes over time, we have the following cases: (a) the Pathogen-ID differs between any of the three cycles, meaning that the read is assigned to different pathogens in one window, (b) the AS/MAPQ score of a read is below a threshold in any of the cycles, and it has been excluded from the analysis in the read assignment procedure. These reads will be considered as false.

The read assignments are based on incomplete fragments of a genome sequence from the intermediate cycles; therefore, the assignments can change during real-time sequence analysis. Thus, the reads can have the same genus assignment in the intermediate cycles but a different assignment in the final cycle. To rectify this issue, (meta-)information from the ground truth cycle is incorporated into the training procedure by substituting the third cycle in each window with the ground truth cycle. During data preprocessing steps, we observed that reads with true observations are frequent and constitute the majority class and false observations are in the minority. Hence, our data consists of imbalanced classes and may lead the model to biased predictions. To overcome this challenge and adjust the class distribution, we used *RandomOverSampler* from Imbalanced-learn [Lemaître *et al*., 2017], which relies on scikit-learn to handle classification with imbalanced classes. It randomly over-samples the minority class (false class in our case) with replacement to create a balanced dataset. The over-sampling is applied only on train data to prevent biased classification and is not used on the test set.

### 2.3 LSTM Classifier

We used Keras [Chollet *et al*., 2015], a deep learning API written in Python, running on top of the machine learning platform TensorFlow [Abadi *et al*., 2015] for the implementation of the LSTM binary classifier. The main properties of LSTM layers are their capability to maintain the internal state of encoded information while iterating over time steps. The LSTM input shape is a tensor with [batch, time-steps, feature] shape. The recurrent *dropout* layer is used for regularization and to avoid over-fitting. The *dense* layer is used for classification. The LSTM built-in layers from Keras used in our model architecture are *keras.layers.LSTM*, with arguments *return-sequences*=*True* in the middle layer. The network architecture contains LSTM built-in layers: one input layer, three layers of LSTM, one dropout layer, and one dense layer. The dense layer uses “*sigmoid*” activation function with “*adam*” optimizer. All the models are trained on 80% of the data and validation on 10%. The remaining 10% is used as a test set. The model is trained in 100 epochs with a batch-size of 200. During the training, *shuffle*=*True* for shuffling the training set before each epoch.

We iterated the modeling by sliding the window for one cycle and evaluated binary prediction performances on the test set. The models’ performances are monitored over time to recognize the best-performing window in any intermediate stages. Ideally, if it is possible to predict the final cycle accurately from intermediate cycles, this can be interpreted as an indication of a possible earlier decision-making time for diagnosis. For this purpose, we defined a criterion such that if the F1 score reaches above 98% in the test set, the predictions are considered sufficiently accurate.

### 2.4 Interpretablility: SHAP

Understanding the reasons behind a prediction is fundamental in assessing any model. We used the SHAP [Lundberg *et al*., 2017] framework to explain predictions of the LSTM binary classifier. This interpretable machine learning paradigm assesses the model locally around the predictions. We used SHAP library’s *Deep.Explainer* for (a) the interpretability of one observation by acquiring the impact of each feature on the prediction output and (b) the distribution of the impacts of each feature on the model output. We conducted experiments on the output of single observations and global model selected from the train and test sets to comprehend the features’ impact on the classification output (true/false). We explain our LSTM classifier concerning the conceptualized binary classifier and the influence of variations of underlying read alignment quality information over time on learning.

### 2.5 Dataset

The dataset used for the evaluation of the method contains the sequencing data of five clinical sputum samples. The QIAamp DNA Microbiome Kit (QIAGEN GmbH) was used for DNA extraction. The extracted DNA samples were amplified via PCR. Library preparation was performed with the Lotus DNA Library Prep Kit (Integrated DNA Technologies, Inc.) using custom sequencing adapters for real-time sequencing. Sequencing was done using an Illumina MiSeq sequencing device with single-end reads of length 151bp (plus 6bp Illumina barcode sequence). For all steps of sample preparation, the original protocols provided by the manufacturers have been followed without adaptions. Real-time analysis of the data was performed based on the algorithm of HiLive2 [Loka *et al*., 2019]. The unprocessed sequencing data is available at the NCBI Sequence Read Archive (SRA) under BioProject number PRJNA825820 (https://www.ncbi.nlm.nih.gov/bioproject/PRJNA825820; human reads have been filtered before data upload). The alignment results serving as input for the presented method are available on Zenodo (https://doi.org/10.5281/zenodo.7547808). The alignment score is contained in the alignment output files using the AS:i tag. The genus is contained in the alignment output files using the GE:Z tag, describing the genus of the reference genome the read was aligned to.

## 3 Results

### 3.1 Classification Results

The LSTM classifier is trained iteratively over eight windows and tested for predicting the final cycle. The prediction performances are monitored for each iteration by evaluating *precision, recall, F1* metrics,

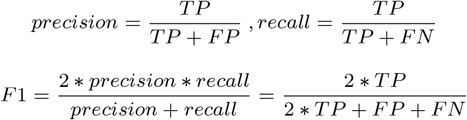

where true positive (TP) refers to the model’s outcomes classified correctly as true, false positive (FP) refers to true outcomes that are classified inaccurately and should have been classified as false, and false negative (FN) refers to false outcomes that should have been classified as true. Classification performances are evaluated for eight windows as shown in Figure 3 to investigate the possibility of accurately predicting final cycle results from intermediate cycles. We observe that accuracy and F1 metrics have an upward trend and a slight growth in window 75-85-95. In this window, the F1 reaches above 98% on the test set and can be interpreted as a highly accurate learned model for the estimation of the final cycle.

**Fig. 3.**
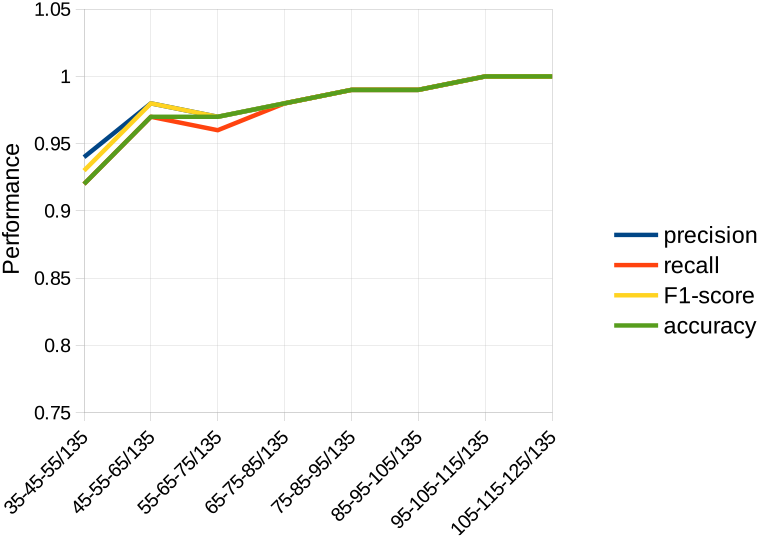
The performances of the trained LSTM binary classifier on the test set is monitored on the y-axis over eight windows on the x-axis based on the sliding window schema depicted in Figure 2. The sequential input is collected from every tenth NGS cycle. In window 75-85-95, F1 reaches above 98% on the test set. This window is identified as a highly accurate model for the estimation of the final cycle.

We evaluated the binary decision results of the model on the reads classified as true to address the speed-accuracy trade-off. The predicated reads with identified microbes are illustrated in Figure 4 for the same eight windows. In orange, Enterobacter is recognized with over twenty short fragments of genome reads in the test set from the beginning. Citrobacter, Streptococcus, and Rothia are detected from the first window with over sixty short fragments of genome reads; however, Escherichia and Raoutella are detected only from the middle cycles onwards. This means that some microbes are detected early without problems, but some need more sequencing time. As we slide the window toward the final cycle, we observe that the predicted number of reads improves over time. However, in window 75-85-95, the same window that has been detected with high accuracy, we have a slight upraise of true reads. This can be associated with a sufficient amount of information available in this window. This time point can be used in detecting the earliest window that matches the ground truth. Overall, the microbes could still be recognized in the sample for this case, significantly before the full length of the read had been acquired. The number of genome fragments also depends on the length of the reference genome; thereby, a high number of reads is not automatically related to stronger detection evidence.

**Fig. 4.**
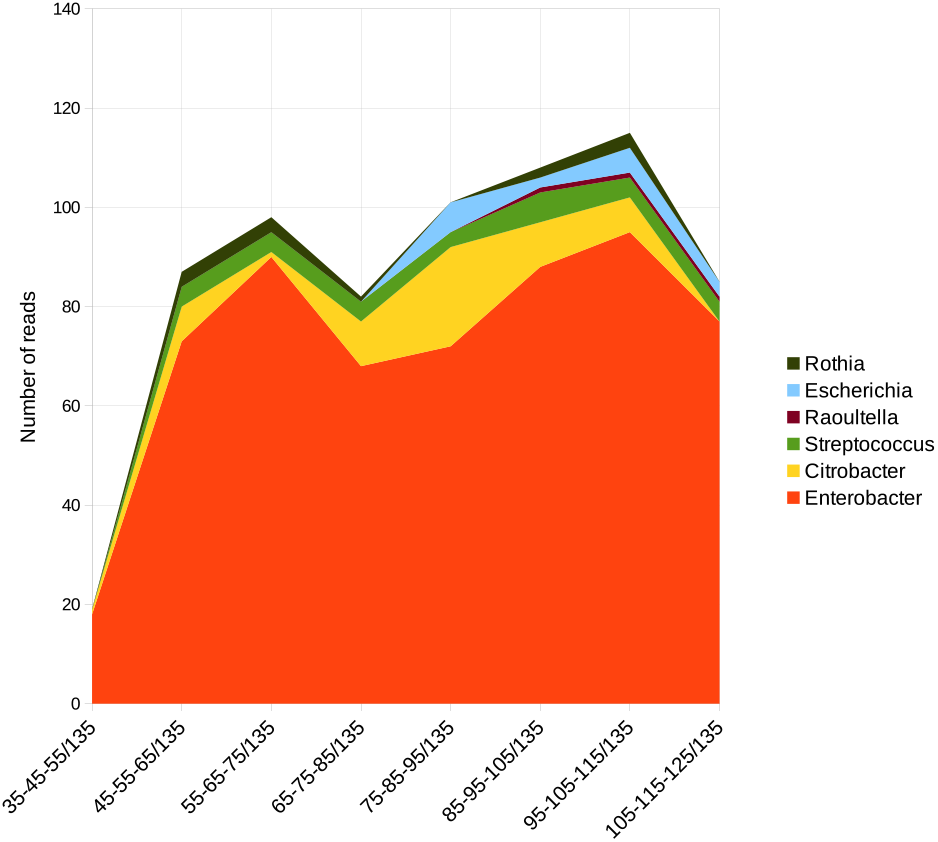
The y-axis is the number of reads identified and matched to microbes, and the x-axis shows eight windows on the test dataset. The genus Enterobacter is recognized with over 20 short fragments of genome reads in the test set from the beginning. Citrobacter, Streptococcus, and Rothia are detected from the first windows; however, Escherichia and Raoutella are only detected from the middle cycles onwards. This means some microbes are detected early without problems, but others need more sequencing time. If the analysis had stopped earlier, Escherichia and Raoutella would have been missing. However, stopping later would lead to the time elapsed without new insights.

The implementations in this research were computationally inexpensive. We could run the preprocessing, training, and testing on the CPU of a PC: 8^th^ Generation Intel Core i7-8665U processor, 1.90 GHz up to 4.80 GHz with Turbo Boost, 4 Cores, 8 Threads, 8 MB Cache, on Ubuntu operating system version 20.04.

### 3.2 Predicting the Decision-Making Time

Our aim in this research is to investigate whether a time point could be predicted to push the decision-making for diagnosis into earlier stages while the real-time NGS analyzer is still running. We applied LSTM as a modeling approach for analyzing read alignment sequential (meta-)information obtained from NGS intermediate cycles.

As we slid the window and trained the LSTM binary classifier on quality score information, we evaluated the model performances to estimate the best-performing model. In window 75-85-95, F1 exceeds our defined criterion of 98% on the test set. If we assume this time point is sufficiently accurate due to capturing the underlying pattern of read alignment (meta-)information, the trained model on this window is expected to make accurate predictions on other samples. We assessed our best-performing model based on this idea. In Figure 5, the F1-scores of the best-performing trained model are evaluated on four different patient datasets on the y-axis and depicted over eight windows on the x-axis on samples 1, 2, 3, and 4. The fifth sample is only used for training and testing models. The F1 is above 75% for all the samples from first window 35-45-55 and reaches 96-98% at window 65-75-85, followed by a dip on window 75-85-95. This can be explained according to our binary classification concept: a read that has been classified as true in an intermediate window is less likely to change alignments in future cycles. However, due to possible seeding cycles of the HiLive2 real-time alignment algorithm in cycles 85 to 95, some read assignments alter during this cycle and/or are discarded during real-time analysis. Another reason can be the completeness of collected information during this window. This means that the assignment procedure in this window is consistent, and most of the reads assigned to microbes are the same as those in the final cycle. On the one hand, if the analysis was stopped in the high-performance window, there would be a possibility of missing the reads assigned to different microbes later. On the other hand, waiting for later cycles will lead to the time elapsed without new insights. Consequently, cycles 85-95 can be associated with the earliest decision-making time, since the F1 reaches above 94% afterwards. Although we have high prediction performances from the beginning of the machine learning analysis, we do not decide based on the very early cycles. As shown in Figure 4, some microbes are detected in early windows, but others need more sequencing cycles; for instance, Eescheria and Raoutella are detected later in this sample. Consequently, it is inefficient to establish the decision-making time on the very early cycles; however, postponing the decision to the end of sequencing will lead to the unnecessarily elapsed time for diagnosis. Overall, still, the microbes could be recognized in the sample for this case before the full-length read was completely acquired.

**Fig. 5.**
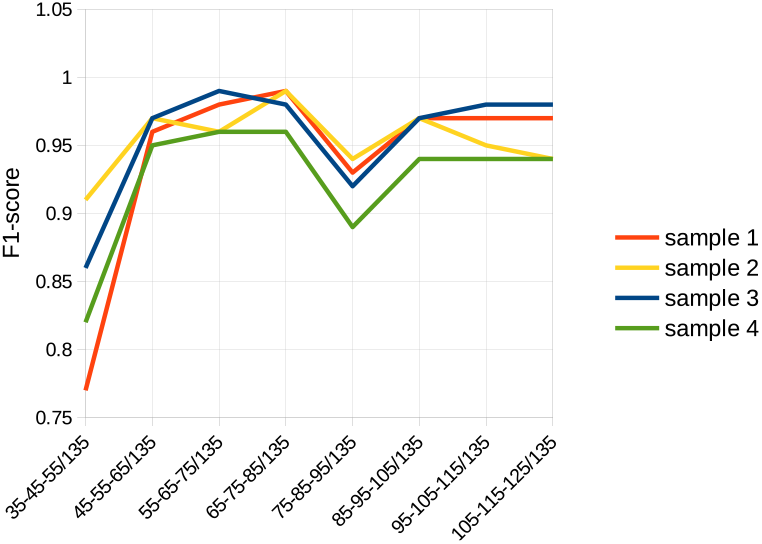
The best-performing LSTM classifier is selected according to the analysis results shown in Figure 3. This model’s performance (F1-score) is then evaluated on four different patient datasets on the y-axis and depicted over eight windows on the x-axis for samples 1, 2, 3, and 4. The F1 is above 75% for all the samples from first window 35-45-55 and reaches 96-98% at window 65-75-85, followed by a dip on window 75-85-95. Due to possible seeding cycles of the HiLive2 real-time alignment algorithm in cycles 85 to 95, some read assignments alter during this cycle and/or are discarded during real-time analysis. If the analysis was stopped in the high-performance window, there would be a possibility of missing the reads assigned to different microbes later. However, waiting for later cycles will lead to the time elapsed without new insights. Consequently, cycles 85-95 can be associated with the earliest decision-making time.

The results show that we can save time to diagnosis by shifting the decision to intermediate cycles observed in the 75-85-95 window, which saves 50 cycles out of 135. Each cycle takes 5 minutes to finish; by this prediction, at least 4 hours could be saved for diagnosis. At the same time, the required sequencing depth and read length are hard to adjust upfront. The proposed method can provide estimation of earliest time for diagnosis; however, this is only possible during runtime when the current alignment results are available. As such, it makes sense to plan a sequencing run with longer reads to validate the intermediate stages.

### 3.3 Explaining the Predictions of LSTM Binary Classifier

Shapley values are means to understand the black-box nature of neural networks. To take action based on a prediction, the interpretability of a machine learning model is essential for decision-makers by assessing trust based on a single prediction and evaluation of a global model to ensure its reasonable behavior [Ribeiro *et al*., 2016]. To incorporate interpretability into our work, we assessed the LSTM classifier by (1) interpreting observations from true and false classes and (2) a summary plot to demonstrate the impact of features on the entire model. First, the classification output of LSTM is assessed for two observations: a read classified as true and another read classified as false to understand the impact of the features on the binary decisions. Additionally, we can interpret how the model differentiates true and false results based on feature values. For a single observation *FOrce.plot* of the SHAP framework generates an explanatory diagram according to the model output; for this study, it is true/false. The features’ impacts are represented by arrows where the larger arrow indicates a higher impact and the smaller arrow carries a lower impact on the binary decision. A red arrow shows a feature that increases the model output value, while a blue arrow decreases it.

We have nine features in every window, and the purpose is to explain the impact of each feature on the classification result. The features and their corresponding cycles are explained in Table1. In each window, quality score information of consecutive cycles are abbreviated as follows, MAPQ: mapping quality, PTH-ID: Pathogen-ID, and AS: alignment score. The cycle number is added to the end of each meta-information as an indicator. MAPQ-C1 corresponds to mapping quality and stored as feature 0, PTH-ID-C1 to Pathogen-ID and stored as feature 1, and AS-C1 to alignment score and stored as feature 2 in cycle 1.

**Table 1.** The features in each window and their corresponding abbreviations are explained. In each window, quality score information of consecutive cycles is used as follows, MAPQ: mapping quality, PTH-ID: Pathogen-ID, and AS: alignment score. The cycle number is added to the end of each meta-information as an indicator. MAPQ-C1 corresponds to mapping quality in cycle 1 and stored as feature 0, PTH-ID-C1 to Pathogen-ID and stored as feature 1, and AS-C1 to alignment score and stored as feature 2.

| cycle number | feature number | meta-information | abbreviation |
| --- | --- | --- | --- |
| C1 | 0 | MAPping Quality | MAPQ-C1 |
|  | 1 | PaTHogen-ID | PTH-ID-C1 |
|  | 2 | Alignment Score | AS-C1 |
| C2 | 3 | MAPping Quality | MAPQ-C2 |
|  | 4 | PaTHogen-ID | PTH-ID-C2 |
|  | 5 | Alignment Score | AS-C2 |
| C3 | 6 | MAPping Quality | MAPQ-C3 |
|  | 7 | PaTHogen-ID | PTH-ID-C3 |
|  | 8 | Alignment Score | AS-C3 |

Shown in Figure 7, in blue, we have Shapley values referring to all the features that pushed the classification prediction value in the negative direction, while the Shapley values in red represent the features that shifted it in a positive direction. In this observation, the read is classified as false where MAPQ-C1, MAPQ-C3, and PTH-ID-C3 in blue, pushed the decision toward 0. The reason is all the features in the negative direction are 0, meaning that the read has been discarded in the last cycle, although it was assigned to a microbe in the first cycle with a low MAPQ. Consequently, the classifier has correctly detected this observation and classified it as 0. AS-C1 and MAPQ-C2 pushed the decision in the positive direction; however, their impacts were not high, and they did not significantly influence the binary decision.

**Fig. 6.**
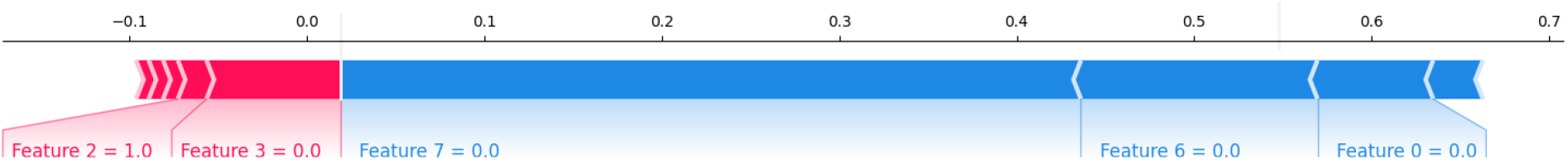
Shapley values are shown for a single observation from the test data classified as false. In blue, we observe the features which pushed the decision toward 0. This read is not assigned to a microbe and has been discarded in cycle 3. Features 0, 6, and 7 correspond to MAPQ-C1, MAPQ-C3, and PTH-ID-C3, in blue, which shifted the decision toward 0. Features 2 and 3 correspond with AS-C1 and MAPQ-C2 pushed the decision in a positive direction; however, they had a low impact on the output.

**Fig. 7.**
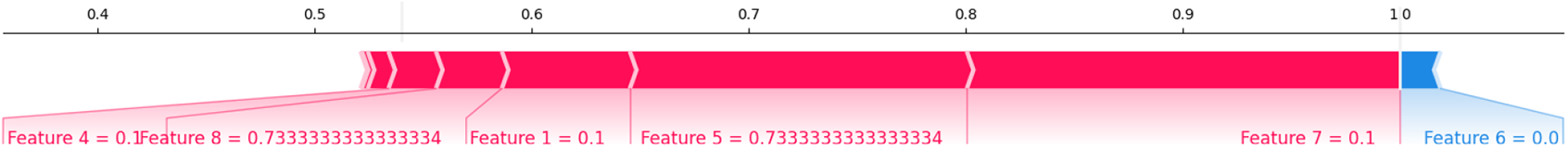
Shapley values are shown for a single observation from the test data classified as true, where the reads continued with the same microbes assignment at all three cycles. The assignment score features 5 and 8 correspond to AS-C2, and AS-C3 are 0.733. The high impact features are PTH-ID-C3, AS-C2, AS-C3, and PTH-ID-C2, features 7, 5, 1, and 8. These features had a positive effect on binary classification results and pushed the binary decision toward 1. MAPQ-C3 is 0, and it had a low impact on pushing the decision in the negative direction.

Figure 8 depicts the Shapley values of an observation classified as true. The features that have positive impacts on the output are PTH-ID-C3, AS-C2, AS-C3, and PTH-ID-C2 and pushed the binary decision toward 1. The AS is 0.733 in cycles 2 and 3, and PTH-ID at all the cycles is 0.1. This means that the read is assigned to the same microbe with high scores, and we expect this read to be classified as true. MAPQ-C3 has a low impact (feature 6 = 0, the lowest possible score) on the output since the read has the same pathogen assignment at all the cycles. Second, to assess the global model, we used *summary.plot* to demonstrate the Shapley values for every feature of every observation on the test dataset. In the summary plot, the features are sorted by the sum of Shapley value magnitudes on all the observations. The plot shows the distribution impact of each feature on the entire output. A high impact value is represented in red, while blue implies the lowering impacts. In Figure 8 Shapley values are shown on the x-axis. All the Shapley values on the left represent the features that shifted the prediction in a negative direction. The points on the right side contribute to shifting the prediction in a positive direction. This summary plot reveals that alignment quality scores PTH-ID-C3, AS-C2, MAPQ-C3, PTh-ID-C1, MAPQ-C2, and AS-C1 have a high impact on classification outcomes as they are sorted by their importance. For instance, we observe lower AS-C2 values have a high negative impact on the prediction value since they spread on the left side, whereas in contrast, higher AS-C2 has a strong positive impact as they spread on the right side. This explanation is valid for MAPQ-C3 and MAPQ-C2. According to alignment scores, high MAPQ/AS scores mean that the read alignments are considered reliable; thus, if a read has higher scores, it is more likely to continue with the same assignment in future cycles. In contrast, if the scores are lower, there are higher chances of discarding the read in the intermediate cycles, and the read is less likely to appear in the final cycle. Thereby, the SHAP analyses of the LSTM classifier confirm the model learned the underlying pattern from (meta-)information obtained from NSG intermediate cycles, and according to our conceptualized binary classification and trained model, which is assessed for single instances and a global understanding of the model.

**Fig. 8.**
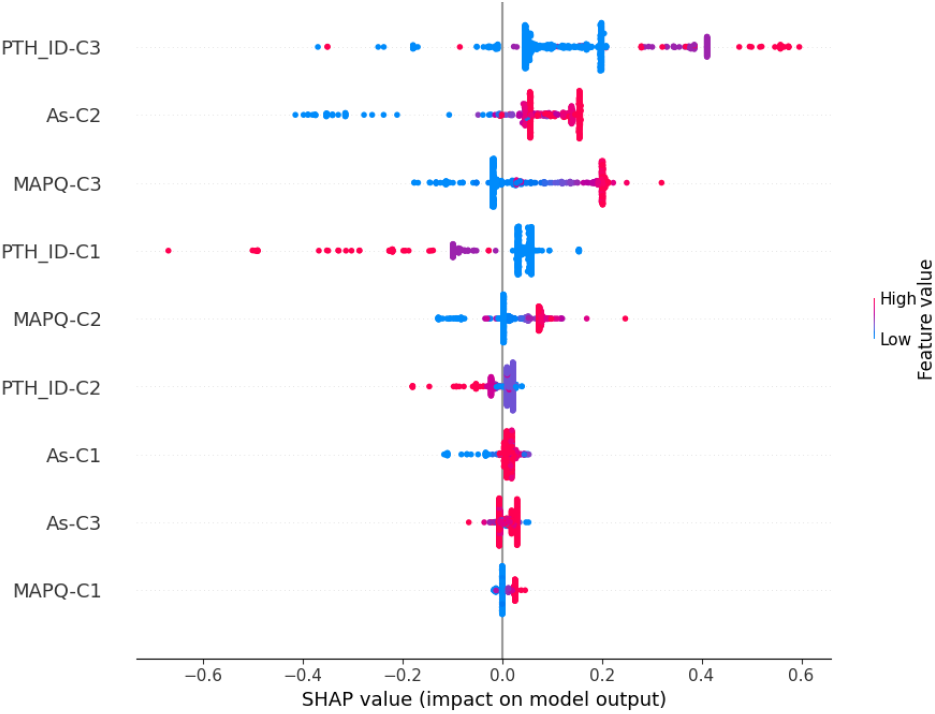
In the summary plot, the features are sorted by the sum of Shapley value magnitudes on all the observations from the test data. It shows the distribution impact of each feature on model output. According to alignment scores, high MAPQ and high AS mean that reads assignment to microbes are considered reliable. Thus, if a read has higher scores, it is more likely to continue with the same assignment in future cycles. In contrast, if the scores are lower, there are higher chances of discarding the reads in intermediate cycles.

## 4 Conclusion

In this paper, we introduced a data-driven approach for analyzing NGS sequencing (meta-)information obtained from intermediate cycles. We proposed a *proof-of-principle* for early diagnosis in time-critical applications. The underlying idea is that by monitoring the predictive performance of trained models on test data and identifying genome reads during intermediate cycles, an approach can be defined for an early time for diagnosis. Monitoring the performance of the LSTM classifier, the best-performing model is selected according to a defined criterion and then tested on different patients’ datasets to comprehend whether the trained model is dependent on a specific dataset. We observed that in intermediate cycles 75-85-95, the predictive performances of the best-performing trained model increased and reached above our criterion and could accurately detect the true genome reads from the final cycle. We can conclude that the LSTM classifier could learn the read alignment quality (meta-)information and applies to different patient samples without retraining the machine learning model or repeating the modeling process. Our experiments verify that our LSTM classifier has high predictive performances on completely unseen data thus, confirms its dataset independent properties. Finally, the intermediate cycle recognized by the best-performing trained model, which has high predictive performance, can be identified as the earliest time point for decision-making. By evaluating the model with the SHAP framework, we confirm that the LSTM classifier is learning the features as defined by our conceptual classification schema, can be trusted, and has the potential for deployment in time-critical diagnosis. We proposed a *proof-of-principle* for predicting early decision-making time in pathogen diagnosis. This approach is based on intermediate results from Illumina sequencing for pathogen diagnosis, but the ideas are more generally applicable in clinical diagnosis and also on other platforms.

## Acknowledgements

We gratefully acknowledge funding by BMBF (Computational Life Science, BMBF - FKZ: 031L0175C) and the Hasso Plattner Institute, Research School on Data Science and Engineering. We acknowledge Suzan Leccese for the generation of libraries used for this study.

